# Prevalence and diversity of TAL effector-like proteins in fungal endosymbiotic *Mycetohabitans spp.*

**DOI:** 10.1101/2023.10.16.562584

**Authors:** Sara C. D. Carpenter, Adam J. Bogdanove, Bhuwan Abbot, Jason E. Stajich, Jessie Uehling, Brian Lovett, Matt T. Kasson, Morgan E. Carter

## Abstract

Endofungal *Mycetohabitans* (formerly *Burkholderia*) spp. rely on a type III secretion system to deliver mostly unidentified effector proteins when colonizing their host fungus, *Rhizopus microsporus.* The one known secreted effector family from *Mycetohabitans* consists of homologs of transcription activator-like (TAL) effectors, which are used by plant pathogenic *Xanthomonas* and *Ralstonia* spp. to activate host genes that promote disease. These ‘*Burkholderia* TAL-like (Btl)’ proteins bind corresponding specific DNA sequences in a predictable manner, but their genomic target(s) and impact on transcription in the fungus are unknown. Recent phenotyping of Btl mutants of two *Mycetohabitans* strains revealed that the single Btl in one *M. endofungorum* strain enhances fungal membrane stress tolerance, while others in a *M. rhizoxinica* strain promote bacterial colonization of the fungus. The phenotypic diversity underscores the need to assess the sequence diversity and, given that sequence diversity translates to DNA targeting specificity, the functional diversity of Btl proteins. Using a dual approach to maximize capture of Btl protein sequences for our analysis, we sequenced and assembled nine *Mycetohabitans* spp. genomes using long-read PacBio technology and also mined available short-read Illumina fungal-bacterial metagenomes. We show that *btl* genes are present across diverse *Mycetohabitans* strains from Mucoromycota fungal hosts yet vary in sequences and predicted DNA binding specificity. Phylogenetic analysis revealed distinct clades of Btl proteins and suggested that *Mycetohabitans* might contain more species than previously recognized. Within our data set, Btl proteins were more conserved across *Mycetohabitans rhizoxinica* strains than across *Mycetohabitans endofungorum*, but there was also evidence of greater overall strain diversity within the latter clade. Overall, the results suggest that Btl proteins contribute to bacterial-fungal symbioses in myriad ways.

**Impact Statement:** Many Mucoromycota fungi harbor endosymbiotic bacteria, including *Rhizopus* spp. that are food fermenters and pathogens of plants and immunocompromised people. *Rhizopus microsporus* has endofungal *Mycetohabitans* (formerly *Burkholderia*) spp. that deploy proteins related to DNA-binding ‘transcription activator-like’ effectors of plant pathogens, which enter plant nuclei and activate disease susceptibility genes. By sequencing isolated bacteria and mining fungal holobiont sequences, we found Btl proteins in diverse *Mycetohabitans* strains, varying in predicted DNA binding specificity, thus in potential host targets. Btl proteins were more conserved within *M. rhizoxinica*, suggesting distinctions among the two named species. The results in the context of phenotypic differences observed in other studies suggest that Btl proteins contribute to symbiosis in diverse ways, providing insight into effector evolution and arguing for functional characterization of additional Btl proteins to understand establishment and maintenance of these important fungal-bacterial interactions.

## Introduction

Effector proteins are synthesized and secreted by bacterial pathogens and mutualists to manipulate their hosts. Investigation of effector protein families reveals mechanisms for disease, resistance, and mutualism, while assessing their evolutionary patterns provides insight into host jumps and commonly targeted host pathways and processes. While some effectors are found across species, many are orphans or belong to smaller groups. Transcription activator-like (TAL) effectors, first discovered in the pepper and tomato pathogen *Xanthomonas euvesicatoria*, have been found in several plant pathogenic *Xanthomonas* (TALEs) and *Ralstonia* (RipTALs) spp. [1]. TAL effectors act in the host as transcription factors to directly upregulate specific host genes. Flanking a central, repetitive DNA recognition domain in TAL effectors are termini containing a signal for the type III secretion system (T3SS), multiple nuclear localization signals (NLSs), and an activation domain [2]. Each repeat in the central domain is composed of 33-35 amino acids that are highly conserved except for two that vary, termed the repeat variable diresidue (RVD) [3]. The target DNA sequence for a TAL effector can be predicted based on the RVDs present, as each RVD dictates the nucleotide-binding specificity of its repeat in a one-to-one manner [4, 5]. Using *tal* gene knockouts, host RNA-seq, and computational prediction of binding sites, in a number of plant species TAL effectors have been found to activate susceptibility genes that benefit the pathogen, including sugar transporters known as SWEETs, transcription factor genes, and a putative sulfate transporter [2]. Notably, not all plant pathogenic *Xanthomonas* and *Ralstonia* spp. or strains within a species have TAL effectors, while some species have more than twenty encoded in their genome [5].

Outside of plant pathogens, TAL effector-like protein sequences have been identified in the marine metagenome, referred to as marine organism TALE-like, i.e., ‘ MorTL,’ proteins [6], and in endofungal Burkholderiales, referred to as *Burkholderia* TAL-like, i.e., ‘Bat’ [7] or ‘Btl’ [8], proteins. Both MorTL and Btl proteins recognize DNA in the same modular fashion as TAL effectors, and they have attracted some interest alongside TAL effectors as building blocks for DNA-targeting reagents [6, 9]. Complete MorTL protein sequences have not yet been obtained, but some Btl protein sequences have. Compared to TALEs and RipTALs, the N- and C-termini of Btl proteins are much shorter, with no clear activation domain, though there is an N-terminal T3S signal and a single, C-terminal NLS [7, 8]. Also, the repeats of the Btl DNA recognition domain are altogether less conserved and contain some RVDs not yet observed in other TAL effectors [8]. While the taxonomic source of MorTL proteins is unknown, the identification of Btl proteins in endofungal Burkholderiales, now classified as *Mycetohabitans* spp. [10], has opened the door to a better understanding of TAL effector evolution and of bacterial-fungal symbiosis.

First identified in 2005, *Mycetohabitans* (formerly *Burkholderia*) *rhizoxinica* (Mrh) is a facultative endosymbiont of *Rhizopus microsporus,* a Mucoromycota fungus that causes rice seedling blight and is an opportunistic human pathogen [11]. Mrh and the related species *Mycetohabitans endofungorum* (Mef) have been studied for their production of secondary metabolites [12, 13], their control of *R. microsporus* reproduction, both sexually and asexually [13, 14], and the mechanisms of their endofungal lifestyle. The bacteria rely on a T3SS to colonize the fungus, but the effectors they secrete through that pathway are unknown, except for Btl proteins [15]. Mechanistic studies have focused on the type strain *M. rhizoxinica* strain B1 [16] and on *Mycetohabitans* strain B13 [17], based on discovery order and tractability in a laboratory setting, leaving the understanding of overall diversity within *Mycetohabitans* limited [8, 18]. The prevalence of *Mycetohabitans* spp. across *Rhizopus* spp. has been surveyed in a few studies [18, 19], suggesting <20% of fungal isolates have endosymbionts. Accessibility of documented symbiotic isolates, lack of symbiosis documentation in culture collection accession metadata, and the small scale of screening conducted so far restricts the ability to conduct large scale analyses of *Mycetohabitans*.

In earlier work, we functionally characterized the only Btl protein, Btl19-13, from Mef strain B13 [8]. It is not required for symbiosis, but it alters the host transcriptome and increases host tolerance to detergent stress, hinting at an influence on cell membrane composition. A Btl protein from a different strain, Btl18-14, with one fewer repeat than Btl19-13 and some differences in the sequence of RVDs was unable to rescue the *btl19-13* knockout. Adaptation to allelic variation at the target in the two respective host isolates, or altogether different functions, are both seen with *Xanthomonas* TALEs [1]. In that same study [8], we discovered by Southern blot that Btl genes or gene fragments are present, in varying number and genomic context, in each of 14 examined strains of *Mycetohabitans* spp. from a global collection of *Rhizopus* accessions; strains were chosen for their availability from culture collections and whether they were previously reported to have endosymbionts, though some novel interactions were identified [19–21]. Recently, another Btl protein was characterized, Btl21-1 (also referred to as Bat1 and MTAL1) from Mrh strain B1 [7, 22]. The predicted DNA binding specificity of Btl21-1 is distinct from that of Btl19-13. A *btl21-1* mutant strain was less successful at colonization due to fungal septa development that prevented the bacteria from moving within hyphae [22], and other Btl proteins from B1 appear to promote successful reestablishment of the symbiosis needed for sporulation [23]. The Southern blot, the negative results of the Btl19-13 and Btl18-14 rescue experiment, and the sequence and functional differences between Btl19-13 and Btl21-1 support the hypothesis that Btl proteins play varied roles in symbioses between different *Mycetohabitans* and *Rhizopus* strains.

Importantly, the distribution and diversity of Btl proteins across *Mycetohabitans* is yet unknown, as is their overall relationship as a group to *Xanthomonas* and *Ralstonia* TAL effectors. Determining Btl distribution and diversity across species and strains from multiple hosts or niches is key to understanding the ecological roles and evolution of these proteins. The limited availability of known *Rhizopus* accessions with *Mycetohabitans* symbionts and genomic resources for those accessions has been a barrier to better understanding how Btl proteins and other symbiotic factors are used. So far, only three *Mycetohabitans* genomes have been sequenced deeply enough to fully assemble the *btl* genes [24, 25]. The repetitive nature of TAL effector-family genes makes them a challenge with low coverage or short-read sequencing. Using long-reads, we sequenced and assembled seven new *Mycetohabitans* genomes and re-sequenced two of the three previously sequenced genomes, and examined the encoded Btl protein sequences to determine their distribution, diversity, and evolutionary relationships. To expand this analysis, we also mined deep, short-read holobiont contig assemblies for a large collection of fungal isolates from the ZygoLife project (Joint Genome Sequencing Project doi: 10.46936/10.25585/60001062) [18, 26–28]. Here we report the findings of this two-part approach, 46 Btl proteins and 48 *btl* gene fragments from Mrh and Mef genomes, revealing widespread distribution and substantial variation in sequence and predicted DNA binding specificity, as well as species-specific patterns of conservation.

## Results

### *De novo sequencing of seven and re-sequencing of two* Mycetohabitans *genomes using long-read technology*

In our prior study [8], we examined *btl* gene content in strains B1, B13, and B14. For this purpose, we used complete genome assemblies of B13 and B14 that we generated by sequencing on the PacBio RSII platform, and an assembly for B1 that was publicly available (GenBank accessions PRJEA51915). During our study, the Joint Genome Institute (JGI) deposited assemblies for B13 and B14 (GenBank accessions PRJNA303198 and PRJNA303197); those yielded the same *btl* gene content as our assemblies, so we cited them in our paper. We have since deposited our B13 and B14 assemblies (**Table 1**). Additionally, we sequenced seven other *Mycetohabitans* strains included in our prior Southern blot analysis [8], using the PacBio Sequel I platform. This set of seven genomes includes that of the type strain of Mef, B5, which was previously only available as a contig-level assembly [10]. We obtained complete assemblies for B5 and five others, with BUSCO genome completeness scores [29] ranging from 99.1-99.4%, and a contig-level assembly for one, B46, with a BUSCO score of 98.2% (**Table I).** B46 was underrepresented in our total read set, likely because it was underrepresented in the multiplexed library. While we were completing our analyses, eight Illumina-based *Mycetohabitans* genome assemblies were made publicly available on NCBI, including some for strains that we sequenced; however, those assemblies are contig level, not closed [30].

**Table 1.** *Mycetohabitans* spp. genome assemblies generated in this study ^1^.

| Strain | Genome Size (bp) | Coverage | Contigs | Chromosome (bp) | Plasmid 1 (bp) | Plasmid 2 (bp) | Protein Coding Genes | tRNAs | %GC | NCBI Accession | Fungal Host Accession |
| --- | --- | --- | --- | --- | --- | --- | --- | --- | --- | --- | --- |
| *Mycetohabitans endofungorum* B3 | 3,541,851 | 463x | 2 | 2,686,710 | 855,141 | N/A | 2,771 | 47 | 60.6% | CP062180-CP062181 | CBS 111563 |
| *Mycetohabitans endofungorum* B5 | 3,355,449 | 402x | 2 | 2,592,175 | 763,274 | N/A | 2,676 | 47 | 61.2% | CP062178-CP062179 | CBS 112285 |
| *Mycetohabitans endofungorum* B60 | 3,610,257 | 434x | 3 | 2,736,534 | 716,486 | 157,237 | 2,869 | 48 | 60.8% | CP062168-CP062170 | NRRL 5560 |
| *Mycetohabitans rhizoxinica* B47 | 3,471,072 | 770x | 2 | 2,646,852 | 824,220 | N/A | 2,805 | 47 | 61.1% | CP062173-CP062174 | NRRL 5547 |
| *Mycetohabitans rhizoxinica* B12 | 3,802,698 | 734x | 3 | 2,776,699 | 857,052 | 168,947 | 2,984 | 47 | 60.7% | CP062175-CP062177 | ATCC 52812; CBS 262.28 |
| *Mycetohabitans rhizoxinica* B49 | 3,533,946 | 925x | 2 | 2,710,688 | 823,258 | N/A | 2,815 | 47 | 60.9% | CP062171-CP062172 | ATCC 46352; NRRL 5549 |
| *Mycetohabitans* sp. B46 | 3,293,065 | 139x | 21 | ND | ND | ND | 2,668 | 51 | 61.0% | JADBGL000000000 | NRRL 5546 |
| *Mycetohabitans endofungorum* B13 | 3,550,038 | 136x | 2 | 2,760,408 | 789,630 | N/A | 2,747 | 47 | 60.6% | CP132744-CP132745 | ATCC 52813 |
| *Mycetohabitans endofungorum* B14 | 3,643,928 | 172x | 3 | 2,674,176 | 768,340 | 201,412 | 2,932 | 47 | 60.6% | CP132741-CP132743 | ATCC 52814 |

<sup>1</sup> ND, not determined; N/A, no second plasmid present.

Our six complete assemblies reveal chromosomes ranging from 2,592,175 to 2,776,699 bp, and one or two megaplasmids each. In all six assemblies there is a megaplasmid of 716-857 kb, and in two, B12 and B60, there is also a smaller megaplasmid, 157 and 169 kb, respectively. The larger plasmids are similar to pBRH01 of the reference strain B1, with the large megaplasmids of B12, B47, and B49 being very similar (99% identity over 91-94% of pBRH01, using megablast), and those of B3, B5, and B60 less similar (87-93% identity over 69-72% of the sequence). The smaller megaplasmids of B12 and B60 have some similarity to the smaller megaplasmid of reference strain B1, pBRH02, with that of B12 being more similar (99% identity over 75% of the sequence) than that of B60 (94% identity over 50% of the sequence).

### Genomic relationships and species of the sequenced strains

To determine the diversity of strains and the species represented by the nine genome assemblies we generated, we performed whole genome phylogenetic analysis [31] including in the analysis the Mrh type strain B1 [25] (**Figure 1A**). Among our newly sequenced genomes is the type strain for Mef, B5. All represented strains grouped as Mrh or Mef except for B46. From this limited number of strains, there was no strong association of species with location or substrate from which the host was isolated: Mrh strains appear in fungal isolates from Asia and North America, and Mef strains in isolates from Asia, North America, Africa, and Europe, and both species appear in fungi isolated from soil or plant material, which represent the majority.

**Figure 1.**
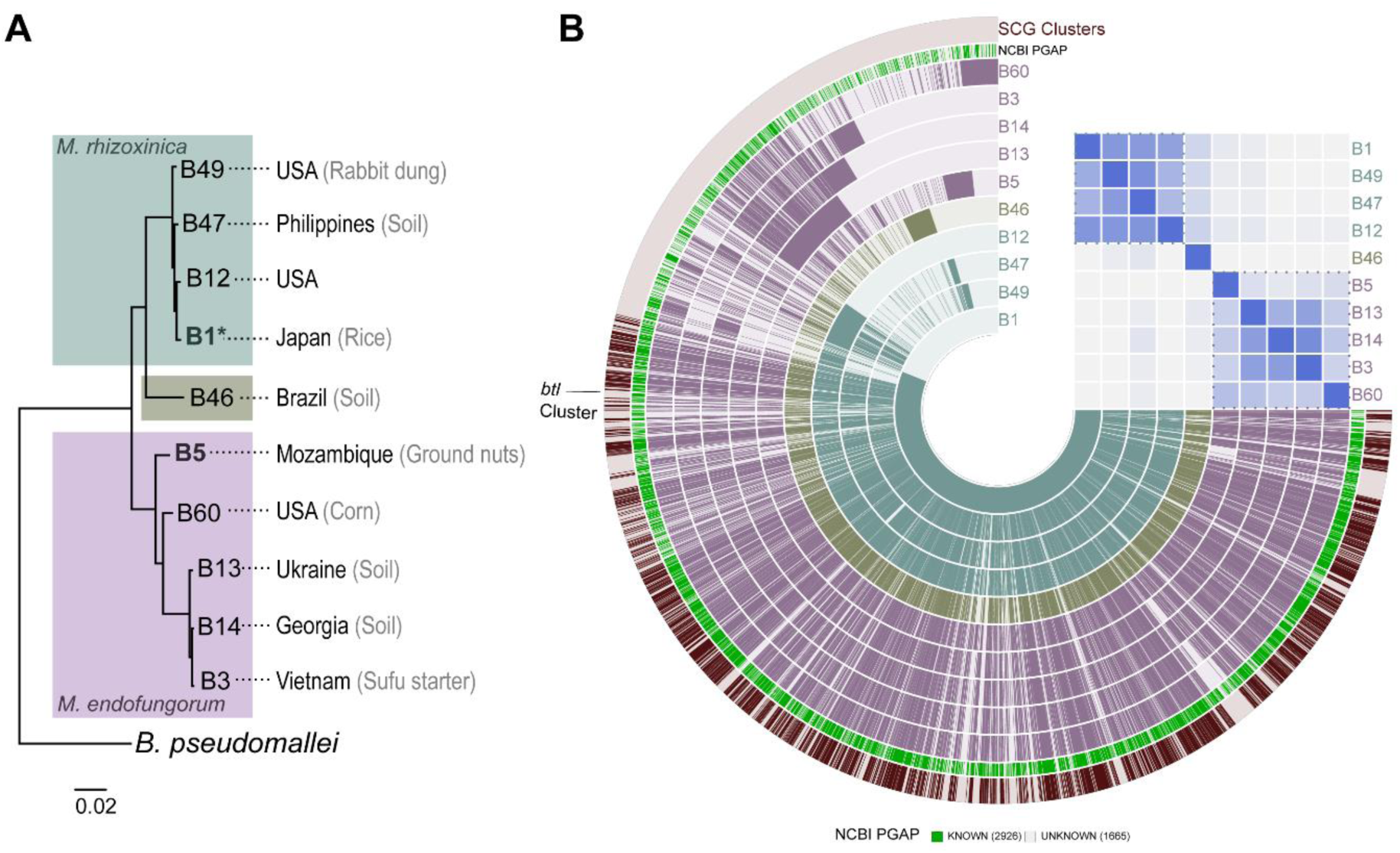
Genomic relationships, species, and origins of the sequenced strains. (A) Output from the reference sequence alignment-based phylogeny builder (REALPHY) for the nine strains sequenced in this study and the *M. rhizoxinica* type strain B1 (denoted by *). B5 is the type strain for *M. endofungorum*. The geographic and substrate origin of the host isolate for each strain is listed when known. *Burkholderia pseudomallei* strain K96243 was used as an outgroup. (B) Pangenome of the ten strains with indicated average nucleotide identity (ANI) heatmap. The blue, green and purple layers represent *M. rhizoxinica, Mycetohabitans* sp. B46, and *M. endofungorum* genomes respectively, where gene clusters are arranged based on synteny with B1. The outermost dark red layer highlights single-copy core gene (SCG) clusters found in all ten genomes. In the heatmap, blue dotted lines indicate clusters of strains with 95% or greater ANI to one another, indicative of species.

We generated a pangenome for the ten *Mycetohabitans* genomes with the help of anvi’o [32]. We used B1 synteny as reference for displaying alignments and included average nucleotide identity analysis, which confirmed our phylogenetic placement of the strains (**Figure 1B**). The pangenome contained 29,051 genes within 4,591 gene clusters. Single-copy core genes (SCGs) were identified in a total of 1,731 gene clusters within the pangenome; all 22 SCGs were found to have 100% annotation based on the Genome Taxonomy Database (GTDB) [33]. We identified 24 putative *btl* genes within the pangenome. These were distributed across all strains except B5 and strains had a maximum of 4 paralogs, found in B1, B12 and B14. These genes were grouped in the same gene cluster and had a combined homogeneity index of 0.73.

### Btl protein diversity across the ten strains

As the goal of our sequencing was specifically to interrogate *btl* genes, we extracted the identified *btl* gene sequences from each of our seven new genome assemblies to compare to the seven known *btl* sequences from the other three genomes [8]. Btl gene sequences were found on the chromosome, the smaller plasmid, the large one, or a combination (**Table S1**). Not all encode full length proteins. Though not identified in our pangenome analysis, a single sequence with homology to *btl* genes could be found in the (complete) assembly for Mef type strain B5 (also called HKI 0456; fungal accession CBS 112285) but it only encodes two repeats, and they are out of frame with each other. Short-read Illumina data for this strain in NCBI (NZ_PRDW00000000.1) confirms this observation, but there is also a deposited sequence (MN891945) encoding a 20-repeat Btl protein attributed to this strain that is not present in our B5 assembly. Similarly, strain B46 from the fungal accession NRRL 5546 contains only a small (479 bp) *btl* gene fragment. The remaining five strains have eleven intact *btl* genes and six *btl* gene fragments between them (**Table S1**). Most of the gene fragments are short, containing one to two repeats at most and many stop codons or frameshifts, and they are often interrupted by transposable elements.

Previously identified full-length *btl* genes have a sequence within ∼200 bp upstream of the start codon that matches the *hrp_II_* box binding element, TTCG-N16-TTCG, for the T3SS regulator HrpB found in *R. solanacearum* and *Burkholderia pseudomallei* [8, 34, 35]. The intact *btl* genes in our newly sequenced strains likewise harbor the element at this location (**Table S2**). Despite this evidence for co-regulation with the T3SS, most of the *btl* genes are not computationally predicted to be substrates for T3S (**Table S2**). Btl19-13 is among these, yet it has been shown to be T3 secreted [8]. Thus, the prediction is lacking, and based on N-terminal sequence similarity to Btl19-13, it seems likely that all the Btl proteins in fact transit the T3 pathway. A possible exception is the eight-repeat Btl protein encoded by a chromosomal gene in strain B60, from fungal accession NRRL 5560 (**Table S2**). This gene lacks the *hrp_II_* element due to an interruption of the putative promoter region and 5’ end of the *btl* ORF by a transposase ORF, which may eliminate expression and is likely in the process of pseudogenization.

To understand the relationships among the different intact Btl protein sequences, we used the QueTAL webtools DisTAL and FuncTAL for phylogenetic analysis [36]. DisTAL removes the RVDs, i.e. the determinants of binding specificity, and aligns based on the remaining repeat ‘backbone’ sequences, while FuncTAL compares only the RVD sequences of the proteins. This allows for differentiation between the potentially rapid evolution of the DNA targeting function versus the presumably slower evolution of the backbone. The Btl proteins group similarly in these two analyses (**Figure 2**). There are differences in the relationships of the clades to each other, and certain proteins, like Btl8-12 and Btl7-1, appear in different relative locations in the two trees. However, the similar clustering overall suggests that the RVDs and the rest of the repeats are evolving in concert. While certain clades comprise only proteins from one *Mycetohabitans* species, such as the set of four 21-repeat Btl proteins present in four Mrh strains, one clade contains sequences from both species. In this small sample set, the degree of conservation varies depending both on the protein and the species.

**Figure 2.**
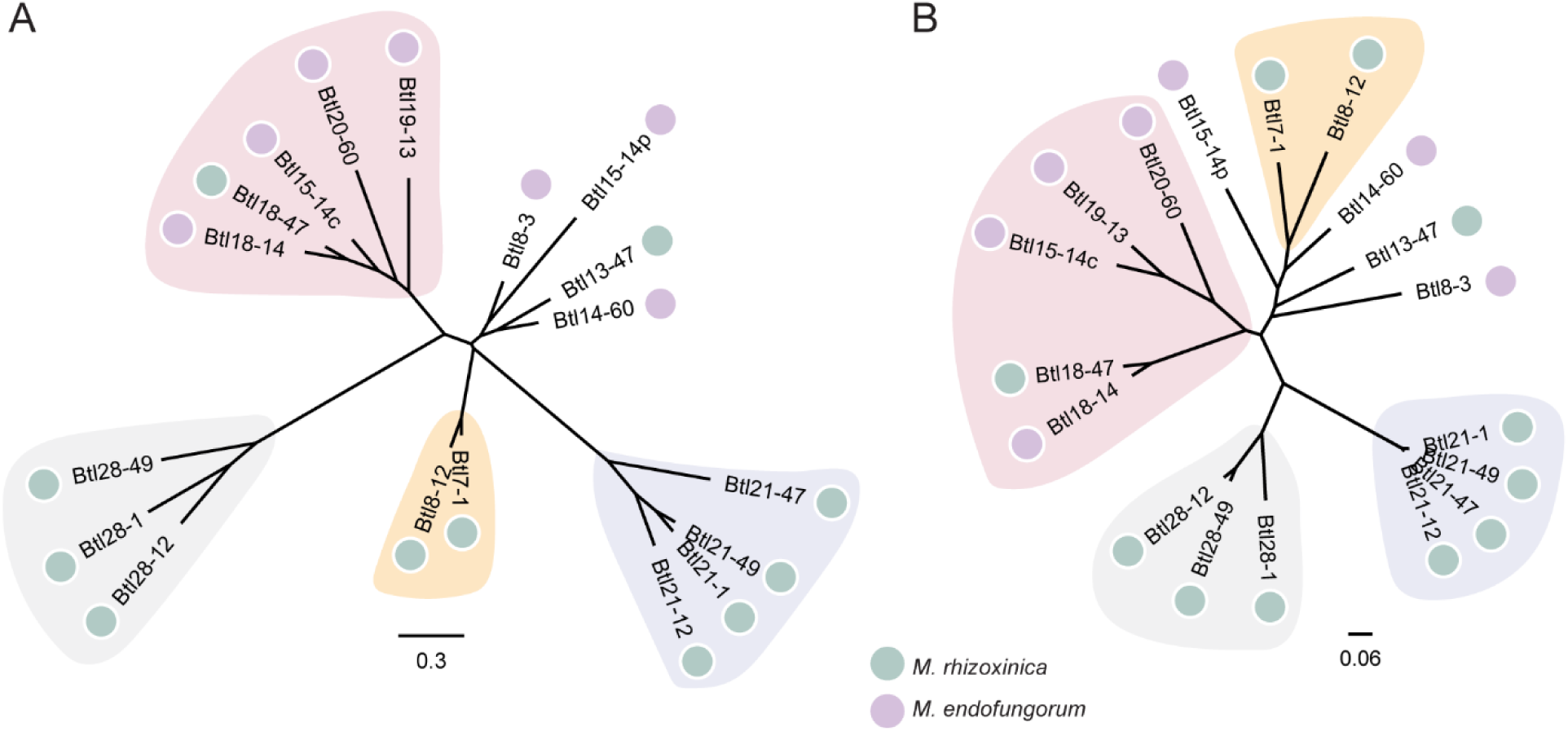
Btl proteins do not exclusively cluster by species. Unrooted trees depicting (A) DisTAL and (B) FuncTAL phylogenetic analyses based on the amino acid sequence with RVDs removed and RVDs only, respectively, for the putative Btl proteins encoded in ten *Mycetohabitans* genomes. More information on these proteins can be found in Table S1. Clades present (with the same members) in both trees are highlighted by background shapes of the same color.

### Btl proteins encoded in fungal hologenomic sequences

To explore Btl protein diversity more fully, we probed genome sequencing data from the ZygoLife Project where low to medium coverage genome sequencing was performed to sample phylogenetic diversity of hundreds of strains in the NRRL collection. The sequences were assembled into metagenomes and contigs that matched bacteria by extracting 16S genes from each assembly and using positive hits to bacteria to prioritize datasets for metagenome-assembled genomes (MAGs). Bacterial MAGs within the data set were previously associated with species using the GTDB Tool Kit [37], however we searched for *btl* sequences in all contigs, not just bacterial assemblies, and indeed identified *btl* gene fragments in two accessions, NRRL 3373 and NRRL A-26124, that did not have sufficient bacterial reads for Autometa assembly and identification of the symbiont genome. New *btl* genes were named using the last two numbers of the fungal accession ID, or, if not unique, the last three.

In addition to the ZygoLife hologenomes, we also searched GenBank for matches to Btl proteins and found deposited fragments from three *Mycetohabitans* strains, HKI 0404/B8, HKI 0513/B6, and HKI 0512/B2, which are symbionts of fungal isolates CBS 308.87, CBS 261.28, and ATCC 20577, respectively [23]. Subsequently, the contig-level genomes of these strains were also deposited (see genome phylogenetics below). Combined, the total number of *btl* genes or fragments identified increased to 94 (**Table 2**).

**Table 2.**
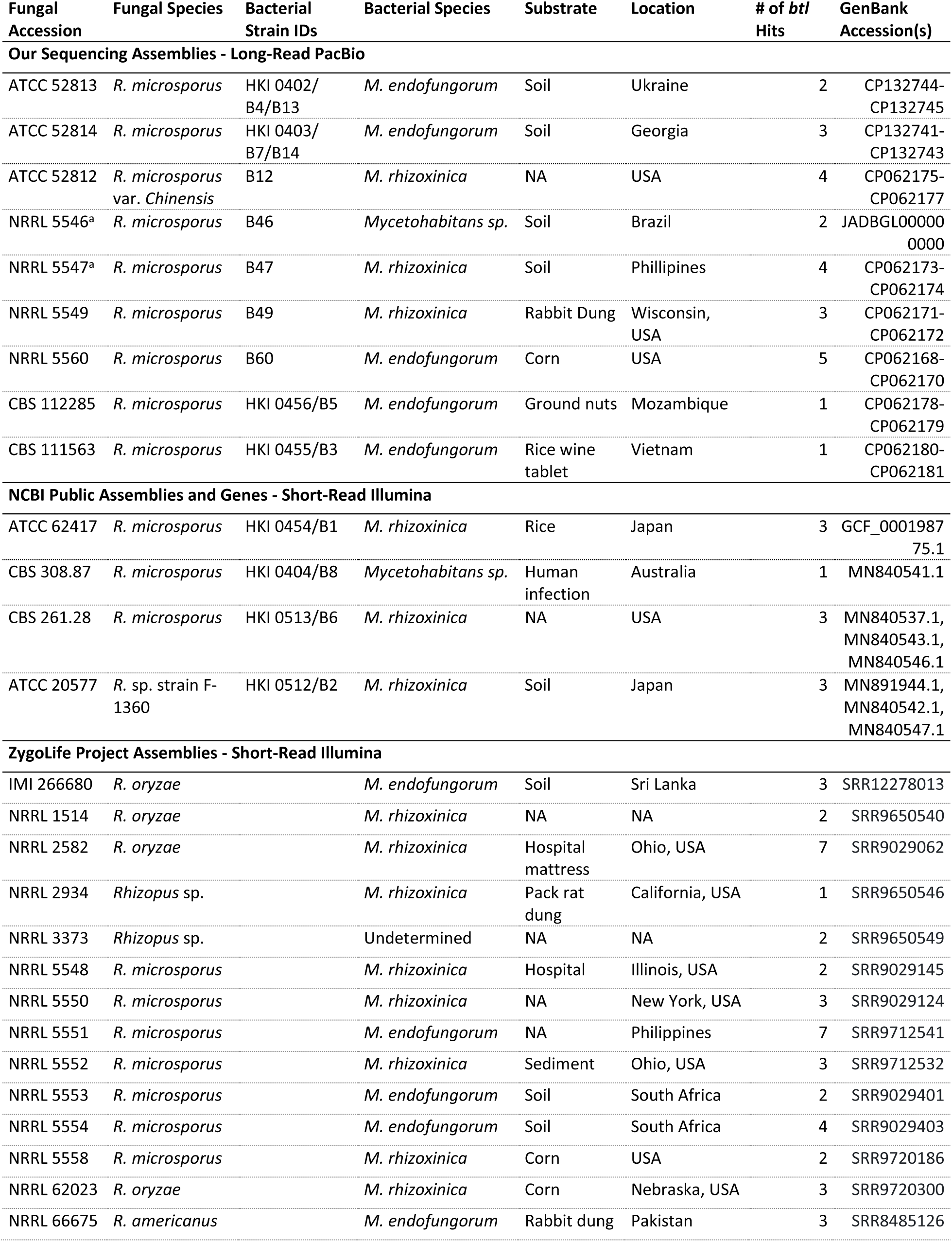
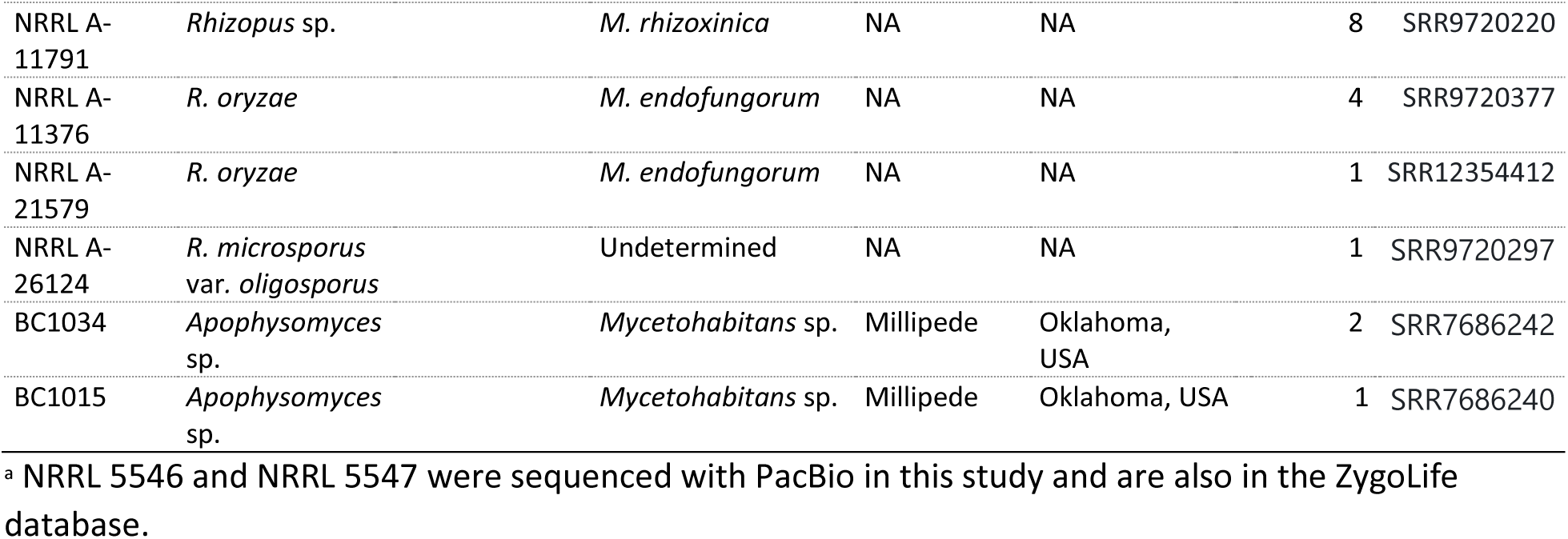
Fungal accessions and their bacterial symbionts examined in this study with the number of *btl* genes or gene fragments detected in each.

Although many *btl* genes within the MAGs fell at the end of a contig and were incomplete, comparison across all of the sequences revealed four mostly conserved types of repeats in Btl proteins (**Figure S1**): (1) a one amino acid shorter pseudo-repeat harboring the RVD HS, HY, NY, or QY; (2) a first, sequence-divergent repeat that has the RVD CD, HD, or YD; (3) the central, standard repeats; and 4) an again sequence divergent final repeat usually harboring the RVD N* (where the asterisk represents a missing residue). This finding agrees with the initial observations of two N- and one C-terminal cryptic repeats in a set of just three Btl proteins analyzed by de Lange et al. [7], who found that the “first repeat” does not confer specificity. Curiously, among other differences, the pseudo, first, and last repeats most often begin with a leucine, while the first residue of the central repeats is commonly phenylalanine. Across the proteins, the final repeat is the least conserved; in some of the Btl proteins this atypical repeat is absent entirely. Comparison across the proteins also revealed variation in the nuclear localization signal following the repeats, either RIRK or QIRK. **Figure S2** shows a maximum likelihood phylogenetic analysis of all Btl protein sequences (complete and incomplete) along with a schematic illustrating the motif variation observed. We chose maximum likelihood over DisTAL for this analysis because, in contrast to the smaller set examined initially 1) the larger data set is more diverse in repeat structure but DisTAL was optimized for highly conserved *Xanthomonas* TALE repeats, and 2) certain pairs within the data set have no overlap within the alignment and DisTAL is built on the neighbor-joining method.

### Hologenomic sequences vs. long-read bacterial genome assemblies

The Mrh strain associated with accession NRRL 5547, hereafter referred to as strain B47, provides a useful case study for comparing the data from the ZygoLife MAGs with our long-read sequenced genomes. We identified three intact *btl* genes in the complete B47 genome. Five different contigs from the B47 MAG contain *btl* homologous sequences: one contains the full length *btl21-47*, and the four others contain the 5’ or 3’ end of *btl18-47* or *btl13-47*, with these fragments falling at the end or beginning of the contig. Highlighting a limitation of short-read sequencing for assembling repetitive sequences like *btl* genes, for each *btl* gene in the B47 MAG, a few repeats within the repeat region captured are missing. However, the translated sequences of the gene fragments match perfectly with the intact Btl18-47 and Btl13-47 encoded in the long-read assemblies, minus the missing central repeats. B46 (from NRRL 5546) was also present in both data sets, but we did not retrieve either of the *btl* fragments in our search of the ZygoLife MAGs.

### *Btl protein diversity in* M. rhizoxinica *relative to* M. endofungorum *strains*

Using the *btl* gene sequences identified in our assemblies, the ZygoLife MAGs, and NCBI, we re-assessed Btl protein diversity, as well as distribution. We excluded incomplete *btl* gene sequences, which may represent assembly gaps or actual pseudogenes, leaving a total of 46 sequences. Six preliminary clades were evident based on tree topology as well as differentiation by motifs, and these were named in numerical order (**Figure 3**). Clade II contains the protein we characterized previously, Btl19-13 [8], but within the clade the number of repeats per protein is not highly conserved, ranging from 8 to 20. Clade III is similarly variable, and these proteins are on average shorter, containing 9-15 repeats. In contrast, the proteins in Clades I and IV are each highly sequence-conserved, including, with some exceptions, repeat number. Clade V consists of sequences from the ZygoLife project only and includes the only proteins with a ‘CD’ RVD in the first repeat. The proteins in Clade VI are truncated to only 7-11 repeats and all members have variants of all key motifs that are unique to this group.

**Figure 3.**
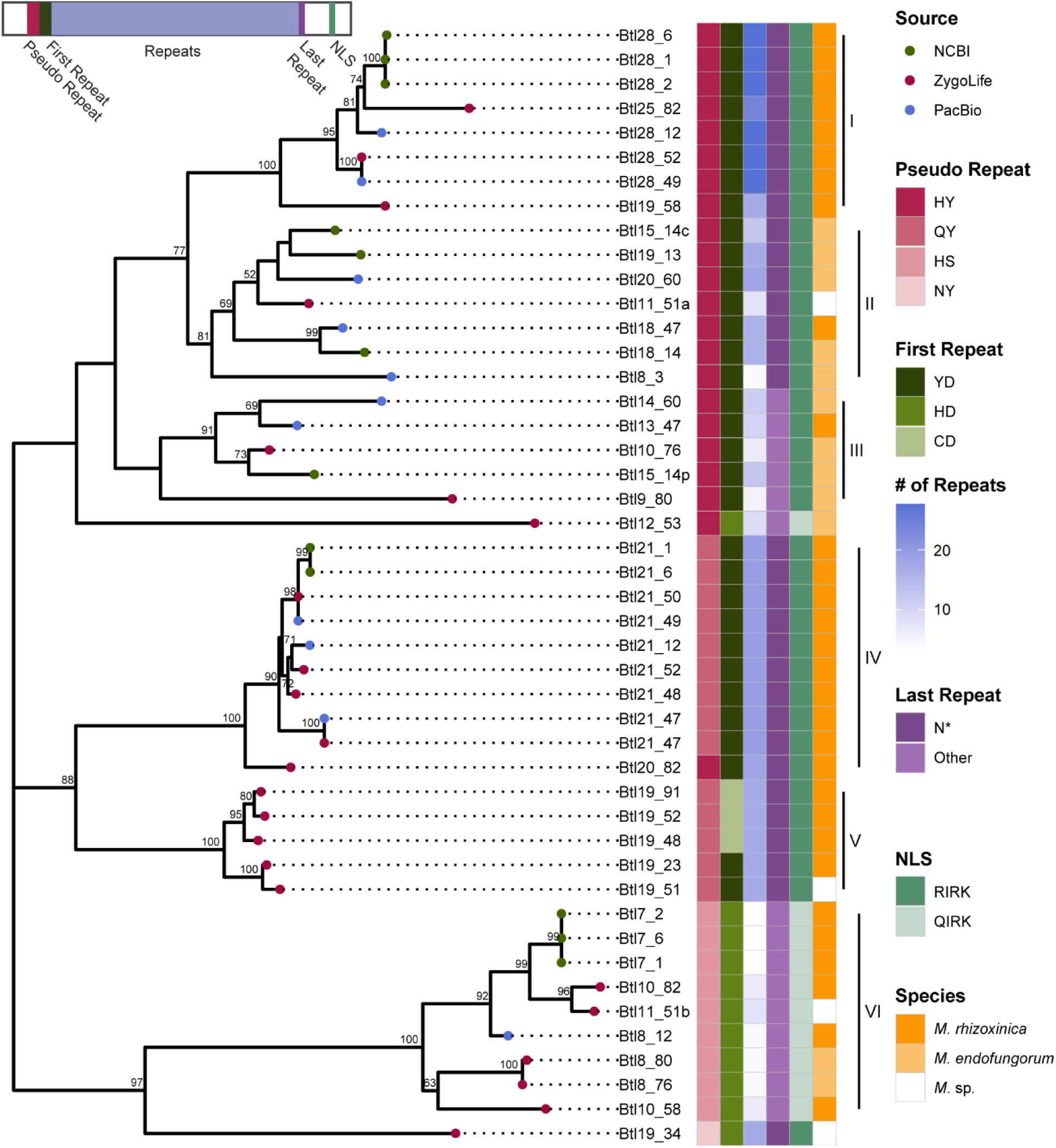
Maximum likelihood phylogenetic analysis of 46 intact Btl protein sequences. Bootstrap values (as % of 1000) are shown at the nodes for those with >50% support. As shown in the key at right, colored circles at branch ends indicate the source of the data for each sequence, and colored boxes detail *Mycetohabitans* species, number of repeats, and presence or absence of key protein motifs, including the nuclear localization signal (NLS), shown using the single letter amino acid code. An asterisk indicates a missing amino acid where one would be expected in a typical conserved repeat, i.e. “N*”. A diagram of a Btl protein with colored, corresponding motif locations is in the top left.

In addition to the whole protein analysis, we extracted and compared just the RVDs of the intact Btl proteins. For automated extraction, we created our own tool (see Methods), which better handles Btl protein repeats than tools designed for *Xanthomonas* TAL effectors. We grouped Btl proteins by clades from **Figure 3** and aligned the RVD sequences to determine if potential DNA targets are conserved within each clade (**Figure 4**). Where there were multiple options for the alignment of an RVD due to a mismatch, deletion, or insertion, we used the backbone repeat sequence to guide the alignment; however, aligned RVDs are not necessarily part of identical repeats. RVD sequences within Clades I, IV, and V are highly conserved within those clades, often only exhibiting RVD polymorphism at only one or two repeats among proteins with similar lengths. In contrast, Clades II and III exhibit relatively high polymorphism protein to protein in the RVDs and in the repeat backbone, even when the RVD is the same. Proteins in Clade VI show an interesting pattern: each but the longest, Btl11-51b, is missing some number of middle repeats relative to that protein. The position and number of repeats missing vary.

**Figure 4.**
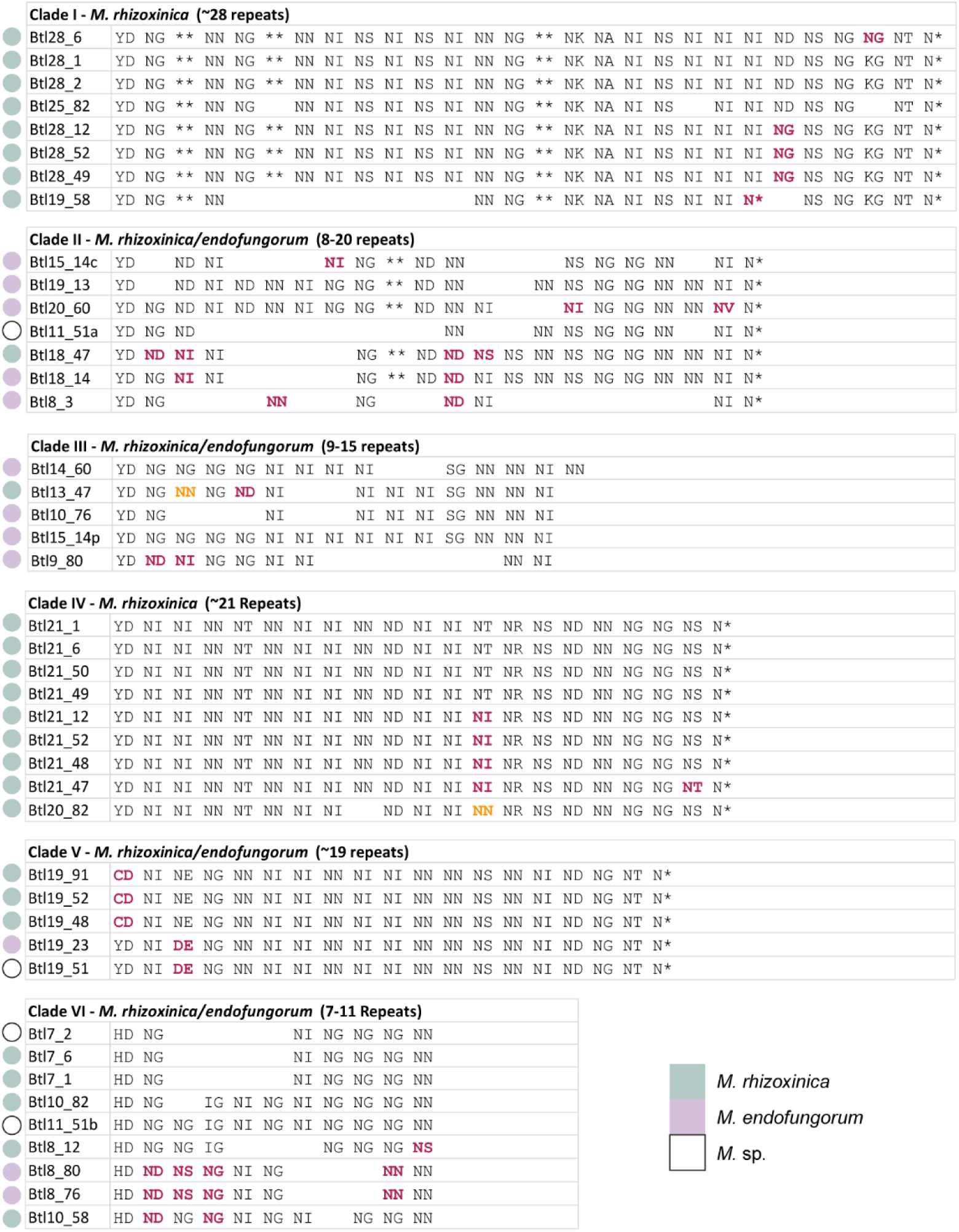
Alignments of Btl protein repeat variable diresidue (RVD) sequences of the first, central and final repeats by clade from Figure 3. RVDs matching the consensus are in black font. RVDs that vary from the consensus are in bold pink font, or bold orange to indicate a second variant at that position. An asterisk (*) represents a missing amino acid at the respective position of the RVD. Colored circles indicate species, following the key at bottom right.

To examine distribution, we carried out whole genome-based phylogenetic analysis on the larger set of strains and mapped the Btl protein content (by clade) onto the resulting tree (**Figure 5a**). We further examined strain relationships by average nucleotide identity (**Figure 5b**). Strains that did not have Autometa bacterial assemblies (NRRL 3373 and NRRL A-26124) or were insufficiently complete (BC1015 and BC1034) were excluded from these analyses. Within this dataset, Mrh strains all have *btl* genes and have more per strain on average than Mef strains. Furthermore, Mrh strains always have a Btl protein from Clade I or VI, and rarely Clades II, III, and V. Both the phylogenetic analysis and the average nucleotide identities suggest that Mef is composed of two species, or possibly subspecies. In one, which contains none of the PacBio-sequenced strains, there are many *btl* gene fragments unassigned to a clade. In the other, which consists mostly of PacBio-sequenced strains, a Btl protein from Clade II is present in all strains, but *btl* gene content varies otherwise. In only one strain, B14 in the latter group, are two Btl proteins from the same clade observed. Two strains reside outside of both groups. The assemblies for these and three strains in the Mef group have only incompletely assembled *btl* genes or pseudogenized *btl* genes or fragments (e.g., B46 and B5).

**Figure 5.**
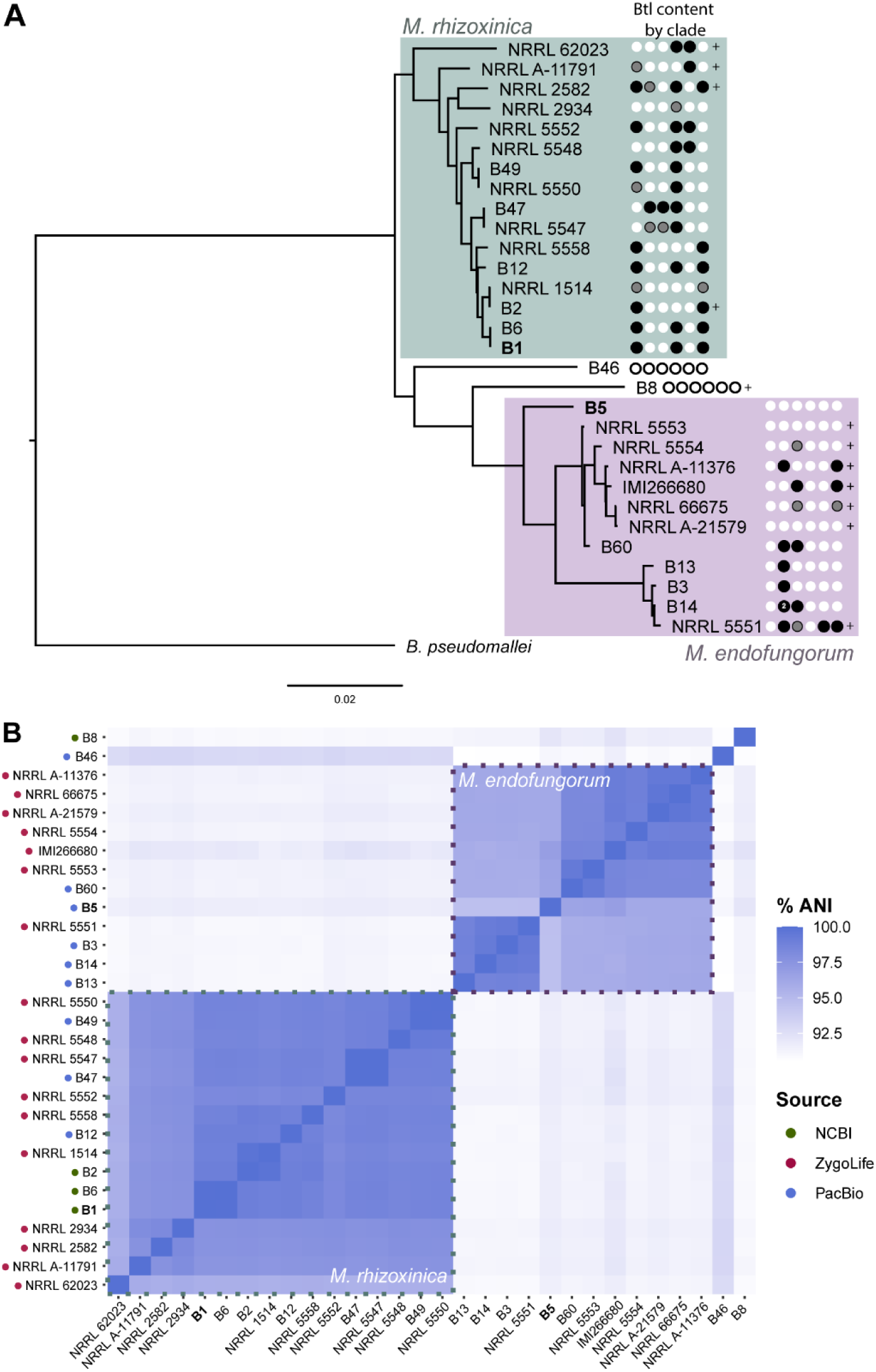
Genomic relationships, species, and *btl* gene content of *Mycetohabitans* strains examined in this study. (A) Output from the reference sequence alignment-based phylogeny builder (REALPHY) for the nine strains we sequenced, the metagenome bacterial assemblies, and publicly available *Mycetohabitans* spp. genomes. Taxa are labeled by bacterial strain name (B#), or fungal accession number where a bacterial strain has not been named. Circles to the right of each taxon represent the *btl* gene content by clade (I-VI from left to right): white represents no gene, black represents one gene from that clade, or two in the case of strain B14, and grey represents an incompletely assembled gene that closely that clade. Plus signs (+) indicate additional incompletely assembled fragments that were not assigned to a clade. *Burkholderia pseudomallei* strain K96243 was used as an outgroup. (B) Average nucleotide identity analysis of the ten strains. Dotted lines indicate clusters of strains with 95% or greater ANI to one another, indicative of species. The source of a strain’s sequence is indicated by colored dots along the left axis. B1 is the type strain for *M. rhizoxinica* and B5 is the type strain for *M. endofungorum*.

### Btl proteins form a distinct but variable clade of TAL effectors

Next, we explored the evolutionary relationship of Btl proteins to TAL effectors and TAL effector-like proteins by generating a maximum likelihood tree based on the amino acid sequences. We chose 1) a representative set of full-length Btl proteins based on **Figure 3, 2**) a set of TALEs from *Xanthomonas oryzae* pv. oryzae (Xoo), *Xanthomonas oryzae* pv. oryzicola (Xoc), *Xanthomonas euvesicatoria* (Xe), *Xanthomonas citri* (Xc), and *Xanthomonas translucens* pv. undulosa (Xtu), 3) RipTALs from *Ralstonia solanacearum* (Rs), 4) all available MorTL sequences [6], and 5) Mycoplasma E-protein sequence fragments, which were suggested to represent distant members of the TAL effector protein family [38]. As shown in **Figure 6A**, Btl proteins cluster distinctly from TAL effectors and RipTALs, which form two closely related clades. One of the MorTL sequences groups more closely with Btl proteins than with the TALE and RipTAL groups. The other appears to have diverged earlier, from a common ancestor of Btls, TALEs, RipTALs, and the first MorTL. The *Mycoplasma* sequences cluster on a separate branch. Finally, to further probe evolutionary relationships, we analyzed the sequences using DisTAL. The more distant of the two MorTL sequences and the *Mycoplasma* sequences were too far diverged from the *Xanthomonas* TALEs to be processed by the algorithm correctly and were therefore not in the resulting tree. However, an interesting pattern emerged (**Figure 6B**): the individual Btl protein branch lengths are on average longer than those of the other proteins in the tree. This difference reflects a greater degeneracy of the repeat backbone sequences within each Btl protein relative to TALEs and RipTALs, hinting at the possibility that they have had more time to diverge or are not as subject as TALEs and RipTALEs to selection for rapid evolution via recombination.

**Figure 6.**
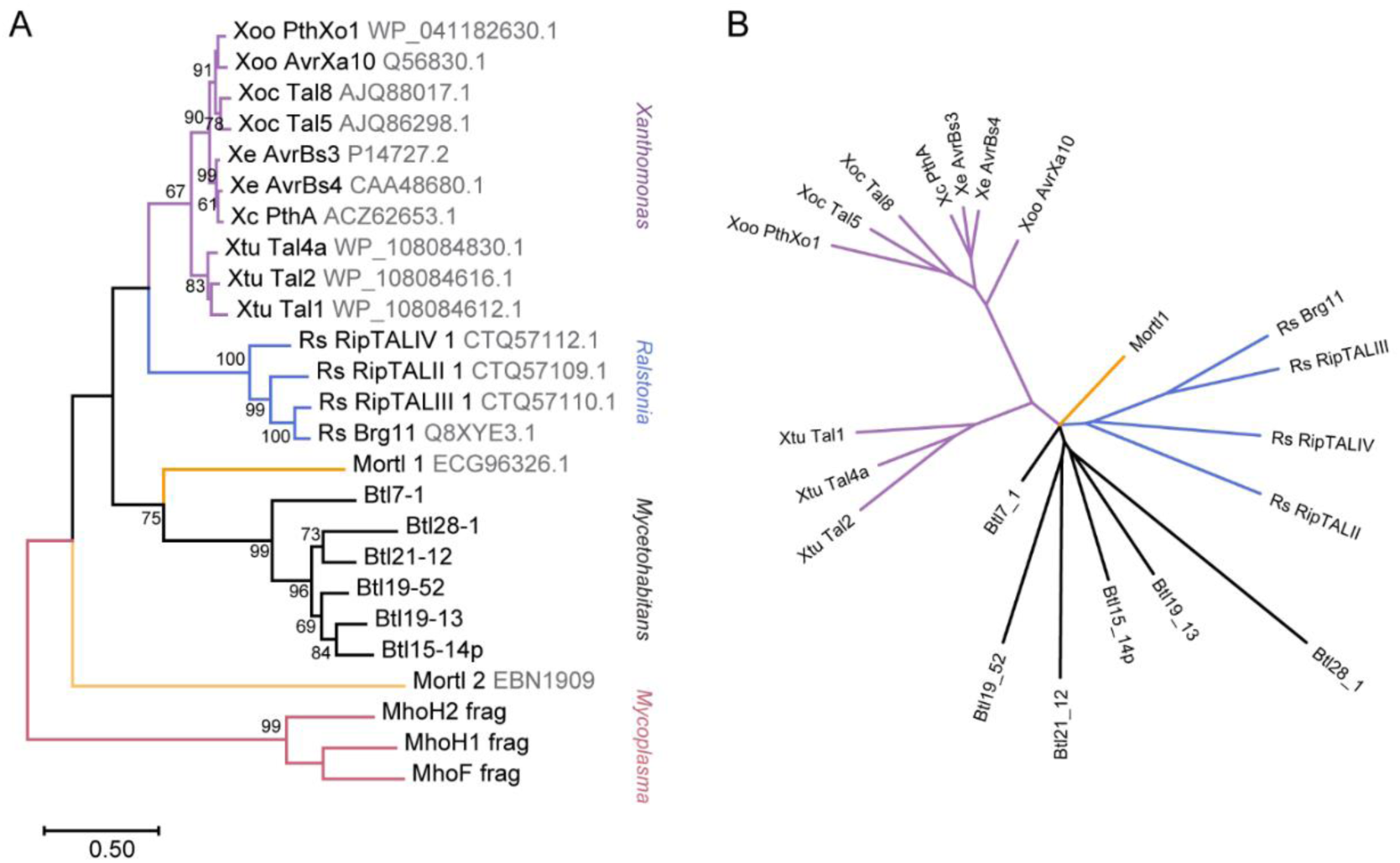
Relationship of Btl proteins to TALEs, RipTALs, and other TAL-like proteins. (A) Maximum likelihood tree based on amino acid sequences of the indicated proteins. Nodes with bootstrap values >50% (out of 1000 replicates) are labeled. The scale represents the number of substitutions per site. Where available, identifiers from UniProt or GenBank are given in gray font next to the protein names. Branches are colored and clades are labeled based on the genus from which the sequence was taken, except for the marine metagenome sequences, which are in orange. (B) DisTAL analysis of the same group of protein sequences presented as an unrooted tree. Line coloring matches that in panel A. Proteins too divergent from *Xanthomonas* TAL effectors to be parsed correctly by DisTAL are absent from the tree.

## Discussion

In this study, we sequenced seven new genomes of *Rhizopus-*associated *Mycetohabitans* spp., six completely and one to a high quality contig-level assembly. We are also making available our complete assemblies for the genomes of two Mef strains that were previously sequenced by the JGI. The *btl* genes extracted from the genome sequences matched prediction by Southern blot [8] and confirmed one to five *btl* genes or fragments per genome, with most genomes having two or three intact *btl* genes, and some having only fragments. Among strains that carry intact *btl* genes, no *btl* gene is conserved across all strains. We expanded our analysis by identifying and retrieving *btl* sequences from fungal metagenome assemblies through the ZygoLife Project. Our ability to extract a large number of intact *btl* genes from the ZygoLife MAGs underscores the value of such large sequencing efforts. Similar to the direct, whole genome sequencing on the smaller set of strains, the metagenome analysis yielded both intact and clearly pseudogenized *btl* genes, as well as incompletely assembled *btl* gene fragments at the ends of contigs. This larger data set enabled analyses that yielded new insight into the conservation and diversification of Btl proteins across *Mycetohabitans* species. The high Btl variability, evidence of psuedogeniziation, and overall lack of conservation across strains are indicative of these effectors undergoing rapid evolution, perhaps bolstered by their repetitive nature.

The number of intact *btl* genes retrieved from the MAGs was somewhat surprising, given the repetitive nature of these genes, the use of short-read sequencing technology in that project, and the anticipated scarcity of bacterial DNA compared to fungal DNA within the library preparations. Indeed, anticipating poor yield and false negatives, we took an inclusive approach to mining: we did not filter metagenomes by symbiont sequence coverage, nor did we filter for *Rhizopus* spp. In total, we identified *btl* sequences in the metagenomes of 19 fungal accessions out of the more than 900 total accessions, of which 200 are *Rhizopus* spp. Interestingly, all accessions yielding *btl* sequences were *Rhizopus* spp., except two, both accessions of a different mucoralean genus, *Apophysomyces* that has also been reported as an opportunistic human pathogen. All of the corresponding bacterial assemblies were identifiable as *Mycetohabitans* spp., except for two, which did not have sufficient bacterial contigs to be extracted as a genome by Autometa during assembly [39].

Importantly, we did not experimentally verify absence of *btl* sequences from accessions for which the metagenome yielded no hits. Yet, we can deduce that some of these are in fact false negatives. Namely, three *Rhizopus* accessions with extracted bacterial genomes had no detected *btl* fragments, yet one of these is B46, which we know from our long-read sequencing assemblies has *btl* gene fragments, albeit short ones. The absence of *btl* gene sequences in the MAGs of the other two accessions may be the result of low overall bacterial sequence coverage. Apart from low coverage, as alluded to earlier, a significant limitation of the MAGs is their having been generated using short-read sequencing.

Compared to the long-read assemblies, short-read assemblies are more prone to fail at the *btl* repeat sequences, leaving it impossible to discern whether a sequence at the resulting contig end is part of a real gene or represents a pseudogene. The short read assemblies and low coverage also result in shorter contigs overall, in some cases yielding *btl* sequences on contigs with few or no other genes, and making it impossible to definitively attribute those sequences to specific bacterial species, or even to the symbiont rather than the fungal host itself. Future, targeted long-read resequencing of some of the hologenomes or isolation and direct sequencing of the bacterial strains would be of interest to fully resolve the *btl* gene repertoires as well as the genomic context for these genes.

One pattern that emerged is a greater conservation of Btl protein content across Mrh than Mef isolates. Each Mrh strain in our analysis has a Btl protein from Clade I or IV, the most conserved Btl clades, with 6 of 10 strains having one of each. Within these conserved clades the RVD sequences are often nearly identical. The observation by Richter et al. [22] that Btl21-1/MTAL1, a member of Clade IV, contributes to the establishment of symbiosis by Mrh strain B1 suggests that Mrh generally may rely on Btl proteins for successful interaction with the fungus. In contrast, the Mef genomes we examined overall have fewer *btl* genes and have no Btl proteins in these conserved clades, and one strain successfully colonizes its host even when its single, Clade II *btl* gene is knocked out [8]. When assessing the diversity of the Mef and Mrh genomes themselves, we do see that Mef strains split into two groups, with all strains in one group containing Btl proteins from Clade II. Yet, those Clade II Btl proteins are still much more variable in repeat sequence than the Clade I and IV Btl proteins found in Mrh strains more distant from one another than the Mef strains are. Altogether, these observations suggest that in contrast to Mrh, Btl proteins play varied roles in Mef, non-essential to symbiosis. The apparent positive effect of Mef Btl19-13 on host membrane stress tolerance [8] however, hints non-exclusively at effects beneficial to the host that may maintain symbiosis over evolutionary time.

It is also possible that some Btl proteins in Mrh and Mef have the same or functionally equivalent host targets, but those targets are sequence-divergent across the Mef host isolates, and the Mef Btl proteins correspondingly adapted and diversified. As Btl proteins have many putative effector binding elements across their hosts genomes, experimental studies validating host targets will be critical for comparison of relevant sequences within host genomes. Or, the Btl protein content variability may be insignificant, simply an artifact of the still relatively small sample size. The Clade V Btl proteins only being from ZygoLife MAGs and some Btl proteins being outliers to the six clades indeed suggests that there are more Btl clades than were captured in this data set.

This study identified many novel Btl proteins, but only a few have been characterized; this is a major caveat to extrapolating Btl function among clade members. Ultimately, functional characterization of representative members of each of the six preliminary clades will help clarify whether the pattern of Btl protein distribution and conservation we observed reflects a diversity of functions, adaptation to a diversity of host genotypes, or some combination, and will lead to a more nuanced understanding of the importance of Btl proteins in *Mycetohabitans-Rhizopus* symbioses. Additionally, though comparative genomics within *Mycetohabitans* spp. analogous to recent studies with endofungal *Mycoavidus* spp. [40] was outside the scope of this study given the limited number of high quality assemblies, identifying and sequencing additional strains would enable a deeper investigation into the *Mycetohabitans* pangenome, shedding more light on the evolutionary trends within and between the species themselves that will inform hypotheses of effector evolution.

## Materials and Methods

### DNA extraction

*Mycetohabitans* spp. were extracted from their fungal hosts by crushing fungal tissue from nonsporulating cultures with a scalpel on an empty petri dish, removing hyphal fragments by syringe filtration, and plating on Lysogeny Broth (LB) amended with 1% glycerol [14]. Plates were incubated at 28°C until colonies formed (approximately 1 week), which were then inoculated to liquid LB and grown with shaking at 28°C until turbid (2-5 days). Genomic DNA was prepared from 3 mL of turbid liquid culture with the MasterPure^TM^ Gram Positive DNA Purification Kit (Lucigen). DNA was quantified by using a Nanodrop^TM^ device (Thermo Scientific) and integrity was assessed by agarose gel electrophoresis.

### Genome sequencing and assembly

Libraries were prepared and sequencing carried out using PacBio long-read technology (Pacific Biosciences), at the Mount Sinai Icahn School of Medicine Genomic Core Facility. B13 and B14 were sequenced on an RSII machine using one SMRT cell each as described [41] and were assembled with HGAP and polished with Quiver in the resequencing pipeline in SMRT Analysis v2.2.0. The seven additional strains were multiplexed and run in the same SMRT cell on a Sequel I machine. Initial assembly was done by the Core Facility using SMRTLink v. 8 (8.0.080529) and the full set of reads. This resulted in closed assemblies with circular contigs for bacterial isolates from accessions CBS112285, CBS111563, and NRRL5549, but several linear contigs each for the others. For the incomplete assemblies, reads were downsampled to 50% using the SMRT Link command line tool bamsieve (using -- percentage 50 and --blacklist options) to create two mutually exclusive, downsampled datasets, and each 50% was separately assembled using SMRTLink v. 9 (9.0.0.92188). This method yielded closed assemblies for genomes of the bacterial isolates from NRRL5547, NRRL5560 and ATCC52812. All closed genomes were run once through the SMRTLink v.9 resequencing pipeline with the complete read set and resulting coverage graphs were inspected for anomalies. Output circular contigs were rotated to the putative replicative origin in SnapGene (www.snapgene.com): chromosomes were rotated to a low GC area just upstream of *dnaA*, the larger megaplasmids to a low GC area immediately upstream of *repA*, and smaller megaplasmids to just upstream of a predicted *repO*. Final assemblies were analyzed for completeness with BUSCO [29].

### Genomic analyses

The Reference sequence alignment based phylogeny builder (REALPHY) pipeline [31] was used to infer the phylogenetic relationships of the ten *Mycetohabitans* genomes using *Burkholderia pseudomallei* strain K96243 as an outgroup (NCBI: BX571965-BX571966). Anvi’o v8 [32] was used to carry out the generation and visualization of a pangenome following the workflow outlined in [42]. Using GenBank files as input, anvi-script-process-genbank was used to generate FASTA as well as NCBI_PGAP-generated gene calls and annotations [43]. Specifically, anvi-gen-contigs-database was run on the FASTA files for each genome with specifying --external-gene-calls as NCBI_PGAP. Functions and gene calls were included in contigs databases using anvi-import-functions. Further, HMMs and SCG taxonomy were run on the contigs databases for further annotation [33, 44]. The pangenome was assembled with the command anvi-pan-genome, utilizing a genomes storage database generated by the command anvi-gen-genomes-storage with the flag for --external-genomes [32].

Additional contig-level genomes from NCBI (B8, GCF_021991875.1; B6, GCF_021991795.1; B2, GCF_021733735.1) and the low coverage strains were processed to produce metagenome-assembled bacterial genomes using SPAdes [45] and binned with Autometa [39] using the pipelines developed for the project (https://github.com/zygolife/DDD). Briefly data were filtered, assembled with SPAdes version 3.15.2 with --meta option and also run in plasmid mode (--meta and –plasmid options) to detect circularized plasmids from metagenomes. Conserved markers and phylogenies were constructed with 16S sequences identified by barrnap v0.9 (https://github.com/tseemann/barrnap). Average nucleotide identity analysis was done using the enveomics collection ANI Matrix tool [46]. Phylogenetic trees were visualized and prepared for publication using FigTree (http://tree.bio.ed.ac.uk/software/figtree/).

### *Identification and annotation of* btl *genes*

In the complete genomes, *btl* gene sequences were identified by querying with the DNA sequence of *btl19-13* using BLAST. The hits were inspected and ORFs annotated manually. For the ZygoLife metagenome assembled genomes (MAGs), a TBLASTN search of non-binned (fungal and bacterial) contigs was performed using the amino acid sequences of Btl19-13 and Btl21-1 as queries, and all contigs with hits to either protein were extracted. A manually curated list of Btl proteins from all available genomes was used to create a reference database in Geneious Prime® (v. 2022.1.1, Biomatters Ltd.), and the “Annotation and Prediction” function was used to annotate the contigs (settings: 50% similarity, “best match” option). Annotation of the contigs with Prokka [v. 1.14.5, 47] and extraction of features containing “effector” from the annotations, and additional tblastn analyses of the contigs using *Xanthomonas* (PthXo1, WP_041182630.1) and *Ralstonia* (RipTALIV-2, LN874063.1) TAL effector sequences as queries were used to validate the final annotations. The promoter region (∼300 bp upstream) of each *btl* gene was inspected for any match to the *hrp_II_* box consensus sequence, TTCG-N16-TTCG [15, 34, 35]. Type III secretion signals were predicted using EffectiveDB [48].

### Phylogenetic analyses of Btl protein sequences

Btl protein sequences from the nine PacBio bacterial assemblies and the reference *M. rhizoxinica* genomes were analyzed using the QueTAL webtools DisTAL (with RVDs removed option) and FuncTAL [36].

Extracted *btl* genes and fragments were translated to amino acid sequences and parsed with an R script called Btl RVD Finder (Github: cartercharlotte/BtlDiversity) to identify key motifs and repeats based on regions conserved among the initial Btl protein set. Btl proteins were named using the number of repeats, starting with the conserved YD repeat as 1, and the bacterial strain number (or abbreviated host strain number when the bacteria are unnamed), following Carter et al. [8]. Proteins lacking a methionine start, a C-terminal NLS, or both the conserved N-terminal pseudorepeat and the YD first repeat were labeled as incomplete. The sequence context of fragments was ascertained manually. Fragments were named in the same way as intact proteins except substituting a unique letter for the (unknown) number of repeats (e.g., BtlA-12 from host strain “12” abbreviated from ATCC 52812).

All amino acid sequences were aligned with MUSCLE and used for a maximum likelihood phylogeny with 1000 bootstraps in MEGAX [49]. The resulting output was converted to a Newick file and visualized using RStudio with the script Btl Phylogeny (Github: cartercharlotte/BtlDiversity). Additional information on species name and data origin was added manually to the table of Btl sequences and features (generated by Btl RVD Finder) and used to annotate the tree. Separately, to eliminate the confounding lack of node support resulting from the presence of the incomplete proteins, a subset of complete Btl protein sequences was similarly processed. Sequence logos were generated [50] from repeat sequences extracted from the final list of all intact Btl proteins.

### All TAL effector family comparison

Representative proteins from all TAL-like families described to date were collected and added to a subset of Btl proteins. From *Ralstonia solanacearum,* four RipTALs were chosen to represent the four designated RipTAL classes [51]: RipTALIV (CTQ57112.1), RipTALII (CTQ57109.1), RipTALIII (CTQ57110.1), and Brg11(Q8XYE3.1). Several *Xanthomonas* TALEs were chosen to represent the known diversity within that family: *Xanthomonas euvesicatoria* AvrBs3 (P14727.2) and AvrBs4 (CAA48680.1), *Xanthomonas oryzae* pv. oryzae PthXo1 (WP_041182630.1) and AvrXa10 (Q56830.1), *Xanthomonas oryzae* pv. oryzicola Tal5 (AJQ86298.1) and Tal8 (AJQ88017.1), *Xanthomonas citri* PthA (ACZ62653.1), *Xanthomonas translucens* pv. undulosa Tal1 (WP_108084612.1), Tal2 (WP_108084616.1), and Tal4a (WP_108084830.1). To represent the family of TAL-like proteins found in the marine metagenome, MOrTL1 (ECG96326.1) and MOrTL2 (EBN1909) [6] were included. Also included were three TAL-like protein fragments identified in a *Mycoplasma* genome [38]. The full set of 25 protein sequences was aligned with MUSCLE and maximum likelihood phylogeny was constructed using all sites with 1000 bootstraps in MEGAX [49]. The same protein set was analyzed with DisTAL [36], though not all proteins were recognized by that software as it was built for *Xanthomonas* TALEs.

## Data Summary

All genomic data are available through the corresponding NCBI accession numbers provided in Table 2. For each of the nine assembled genomes, *btl* genes are annotated within the genome accession. For the metagenome-assembled genomes, *btl* genes and fragments have been deposited separately as a third-party annotation (Accessions: BK063803-BK063866). Fasta files for the Btl protein amino acid sequences and scripts from R analysis are available on GitHub (cartercharlotte/BtlDiversity).

## Acknowledgements

Support for M.E.C. and the single strain sequencing came from the U.S. Department of Agriculture (USDA) National Institute of Food and Agriculture (EWD 2018-67011-28015 and 2021-67034-40327). B.L. was supported by the USDA (USDA-ARS Project 8062-22410-007-000D). Microbial strains used in this work were provided by the USDA-ARS Culture Collection (NRRL). J.E.S. is a CIFAR fellow in the program Fungal Kingdom: Threats and Opportunities and was supported by the National Science Foundation (NSF) for (DEB-1441715 & EF-2125066) and the USDA (National Institute of Food and Agriculture Hatch projects CA-R-PPA-211-5062-H). Analyses were performed on the UC Riverside High Performance Computing Cluster supported by the NSF (DBI-1429826 & DBI-2215705) and the National Institutes of Health (NIH) (S10-OD016290) and the UNC Charlotte Research Cluster. The Zygolife project data was supported by NSF grants DEB-1441604, 1441677, 1441715, 1441728. We also thank the JGI for sequencing produced under proposal 10.46936/10.25585/60001019. JGI (https://ror.org/04xm1d337) is a Department of Energy User Facility supported by the Office of Science of the U.S. Department of Energy operated under Contract No. DE-AC02-05CH11231.

