## Supplemental Materials for "Prevalence and diversity of TAL effector-like proteins in fungal endosymbiotic *Mycetohabitans spp.*"

**Table S1.** *btl* genes and gene fragments (gray background) in ten *Mycetohabitans* spp. genomes.

| Fungal Strain | Bacterial Strain | Gene Name[1] | Replicon | Locus | Coordinates | Translated Protein RVDs |
|---|---|---|---|---|---|---|
| ATCC 62417 | HKI 0454/ B1 | *btl21-1* | Plasmid 1 | RBRH_01844 | 565327...567642 | YD NI NI NN NT NN NI NI NN ND NI NI NT NR NS ND NN NG NG NS N* |
| | | *btl28-1* | Plasmid 2 | RBRH_01770 | 106351...109344 | YD NG ** NN NG ** NN NI NS NI NS NI NN NG ** NK NA NI NS NI NI NI ND NS NG KG NT N* |
| | | *btl7-1* | Plasmid 2 | RBRH_01778 | 104797...105732 | HD NG NI NG NG NG NN |
| ATCC 52813 | HKI 0402/ B4/B13 | *btl19-13* | Chromosome (CP132744) | RA167_03370 | 728644...730788 | YD ND NI ND NN NI NG NG ** ND NN NN NS NG NG NN NN NI N* |
| | | - | Chromosome (CP132744) | RA167_03350 | 723396...723821 | - |
| ATCC 52814 | HKI 0403/ B7/B14 | *btl15-14c* | Chromosome (CP132741) | RA166_09505 | 2089484..2091232 | YD ND NI NI NG ** ND NN NN NS NG NG NN NI N* |
| | | *btl15-14p* | Plasmid 2 (CP132743) | RA166_16070 | 95736..97463 | YD NG NG NG NG NI NI NI NI NI NI SG NN NN NI |
| | | *btl18-14* | Plasmid 2 (CP132743) | RA166_16105 | 106266..108311 | YD NG NI NI NG ** ND ND NI NS NN NS NG NG NN NI N* |
| | | - | Chromosome (CP132741) | RA166_09530 | 2097598..209713 | ND NG NN |
| CBS 111563 | HKI 0455/ B3 | *btl8-3* | Chromosome (CP062181) | IHE27_12345 | 2068391..2069461 | YD NG NN NG ND NI NI N* |
| CBS 112285 | HKI 0456/ B5 | - | Plasmid 1 (CP062179) | IHE28_14160 | 612407..612753 | - |
| ATCC 52812 | B12 | *btl28-12* | Plasmid 2 (CP062177) | IHE29_16130 | 68949..71942 | YD NG ** NN NG ** NN NI NS NI NS NI NN NG ** NK NA NI NS NI NI NI NG NS NG KG NT N* |
| | | *btl21-12* | Plasmid 2 (CP062177) | IHE29_16140 | 72678..74996 | YD NI NI NN NT NN NI NI NN ND NI NI NI NR NS ND NN NG NG NS N* |
| | | *btl8-12* | Plasmid 2 (CP062177) | IHE29_16125 | 67296..68330 | HD NG NG IG NG NG NG NS |
| | | - | Plasmid 2 (CP062177) | IHE29_16110 | 63680..63913 | - |

| | | | | | | |
|---|---|---|---|---|---|---|
| NRRL 5546 | B46 | - | Contig 3 | IHE30_05195 | 280204..280683 | - |
| | | - | Contig 8 | IHE30_10450 | 136858..137049 | - |
| NRRL 5547 | B47 | *btl18-47* | Plasmid 1 (CP062173) | IHE31_00180 | 45803...47848 | YD ND NI NI NG ** ND ND NS NS NN NS NG NG NN NN NI N* |
| | | *btl21-47* | Plasmid 1 (CP062173) | IHE31_02255 | 598438...600753 | YD NI NI NN NT NN NI NI NN ND NI NI NI NR NS ND NN NG NG NT N* |
| | | *btl13-47* | Plasmid 1 (CP062173) | IHE31_00165 | 40437...41966 | YD NG NN NG ND NI NI NI NI SG NN NN NI |
| | | - | Chromosome (CP062174) | No ORF | 638061..641324 | - |
| NRRL 5549 | B49 | *btl28-49* | Chromosome (CP062171) | IHE32_09390 | 2124213..2127206 | YD NG ** NN NG ** NN NI NS NI NS NI NN NG ** NK NA NI NS NI NI NI NG NS NG KG NT N* |
| | | *btl21-49* | Plasmid 1 (CP062172) | IHE32_14660 | 621296..623611 | YD NI NI NN NT NN NI NI NN ND NI NI NT NR NS ND NN NG NG NS N* |
| | | - | Chromosome (CP062171) | No ORF | 590914..591150 | - |
| NRRL 5560 | B60 | *btl20-60* | Plasmid 1 (CP062169) | IHE33_14225 | 506369...508612 | YD NG ND NI ND NN NI NG NG ** ND NN NI NI NG NG NN NN NV N* |
| | | *btl14-60* | Plasmid 1 (CP062169) | IHE33_14240 | 512461...514089 | YD NG NG NG NG NI NI NI NI SG NN NN NI NN |
| | | - | Chromosome (CP062168) | IHE33_09065 | 2121146..2121352 | - |
| | | *btl8-60*[2] | Chromosome (CP062168) | IHE33_09045 | 2117750..2118772 | NI NS NG NG NN NN NV N* |
| | | - | Plasmid 2 (CP062170) | IHE33_15585 | 102460..103052 | - |

[1]*btl* gene names are the repeat number, a dash, then the strain number. Repeat number includes the conserved YD/HD repeat that precedes specificity-defining repeats. The letters p and c distinguish plasmid- vs. chromosomally-located genes in a strain if the genes have the same number of repeats.

[2]Interrupted by a transposase gene insertion at the 5' end.

**Table S2.** Predicted type III secretion signal and putative *hrp$_{II}$* box for full-length *btl* genes from the seven newly sequenced *Mycetohabitans* genomes.

| **btl *genes*** | ***Locus*** | ***Predicted T3SS signal*[a]** | ***hrp$_{II}$ Box Sequence (TTCG-N16-TTCG)*** | ***hrp$_{II}$ box position*** | ***Distance between hrp$_{II}$ box and ATG*** |
|---|---|---|---|---|---|
| *btl8-3* | IHE27_12345 | None | CGAACGCTTCACCTACTGGGCGAA | 2069677..2069700 | 215 |
| *btl28-12* | IHE29_16130 | 0.99902 | TTCGCTCAGTAGGTGAAGCGTTCG | 68710..68733 | 215 |
| *btl21-12* | IHE29_16140 | None | TTCGCCCAGTAGGTGAAGCGTTCG | 72449..72472 | 205 |
| *btl8-12* | IHE29_16125 | None | TTCGCCGGATCGGCGGTGCGTTCG | 67076..67099 | 196 |
| *btl21-8* | IHE29_16125 | None | TTCGCCGGATCGGCGGTGCGTTCG | 67076..67099 | 196 |
| *btl18-5547* | IHE31_00180 | None | TTCGCCCAGTAGGTGAAGCGTTCG | 45564...45587 | 215 |
| *btl21-5547* | IHE31_02255 | None | TTCGTTCAGCACGCAATGCGTTCG | 600960..600983 | 206 |
| *btl13-5547* | IHE31_00165 | None | TTCGCCGGATCGGCGGCGCGTTCG | 40217...40240 | 196 |
| *btl28-5549* | IHE32_09390 | 0.99902 | TTCGCCCAGTAGGTGAAGCGTTCG | 2127422...2127445 | 215 |
| *btl21-5549* | IHE32_14660 | None | TTCGTTCAGCACGCAATGCGTTCG | 623818..623841 | 205 |
| *btl20-5560* | IHE33_14225 | 0.99902 | TTCGCCCAGTAGGTGAAGCGTTCG | 508828..508851 | 215 |
| *btl14-5560* | IHE33_14240 | None | TTCGCCGGATCGGCGGCGCGTTCG | 514286..514309 | 196 |

[a] T3SS signal prediction was carried out using EffectiveDB (36).

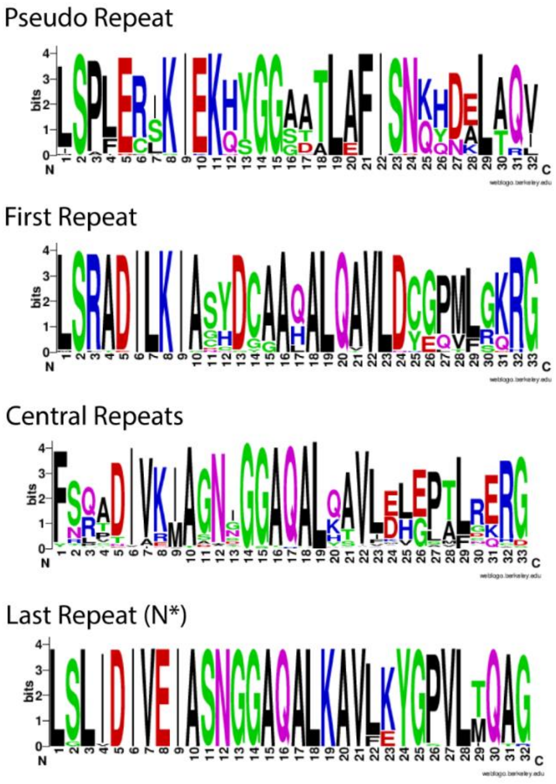


**Figure S1.** Sequence logo motifs for the pseudo repeats, first repeats, central repeats, and terminal repeats of all intact Btl proteins used in Figure 3. For the last repeats, only those with an N* RVD (the asterisk denoting a missing residue) were included, as those with other RVDs are indistinguishable from central repeats.

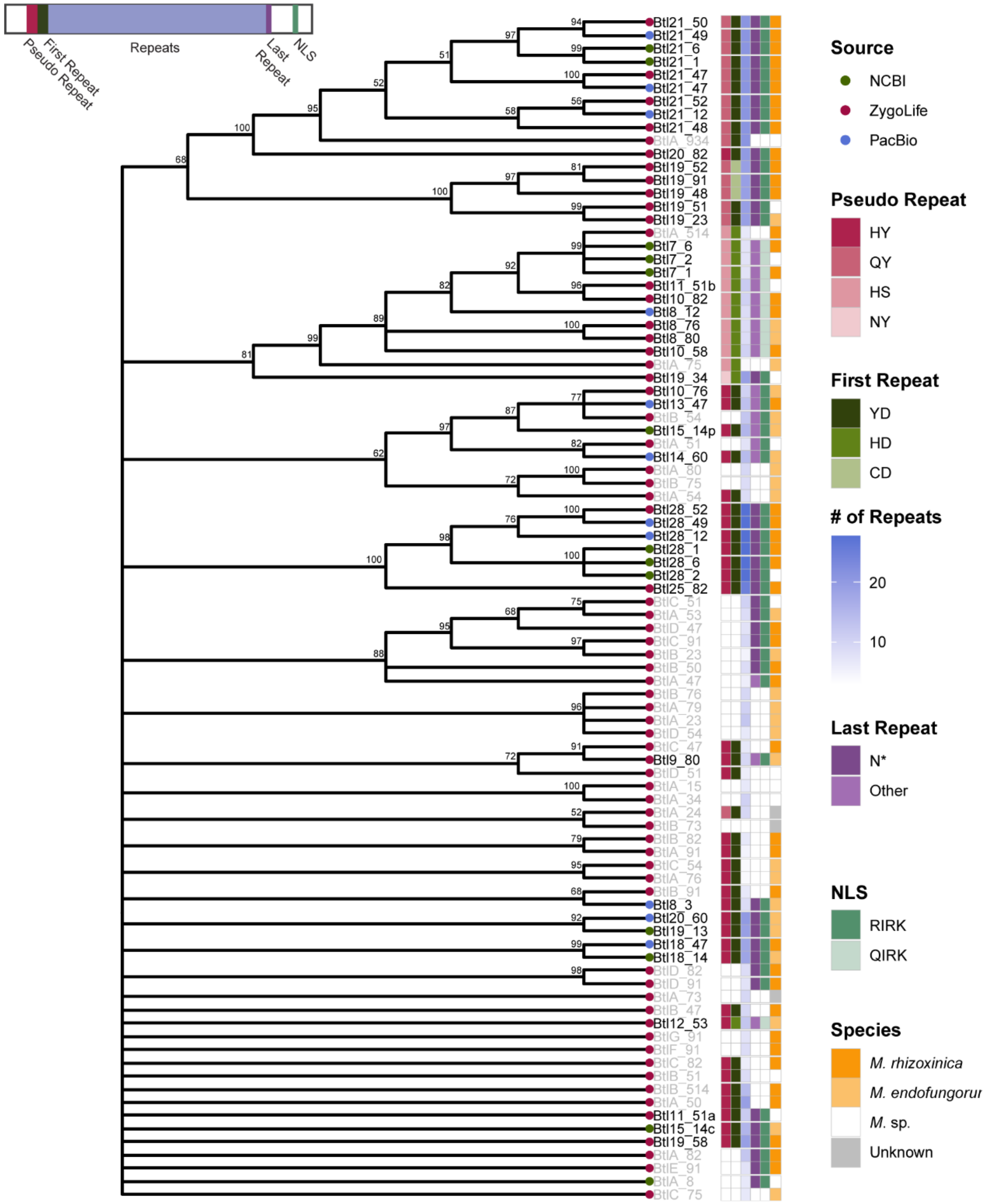


**Figure S2.** Maximum likelihood phylogenetic analysis of encoded Btl protein sequences across 32 isolates. Grey tip labels indicate incomplete protein sequences, typically found encoded at the end of an assembled contig; these are named using a letter in place of the repeat number. Branches with bootstrap values less than 50% were collapsed; those with >50% are labeled at the nodes. As shown in the keys on the right, colored circles at tips indicate the source of the data for each sequence, and colored boxes detail *Mycetohabitans* species, number of repeats, and presence or absence of key protein motifs, including the nuclear localization signal (NLS), shown using the single letter amino acid code. A diagram of a Btl protein with motif locations is in the top left.
